# Integrating glycosylation in *de novo* protein design with ReGlyco Binder Design Filter

**DOI:** 10.64898/2026.04.16.718906

**Authors:** Ojas Singh, Elisa Fadda

## Abstract

Artificial Intelligence (AI)-based methods for 3D protein structure prediction are revolutionising structural biology, rapidly giving users a 3D perspective on any molecular architecture and protein-protein interaction (PPI) virtually on-demand. Regardless of the inherent limitations of the various methods to date, the continuous improvement of the algorithms, and the broad availability of open access (OA) web servers, software packages and databases are bound to accelerate biopharmaceuticals’ discovery. Within this context, the development of computational pipelines for the *de novo* design of target-specific protein binders is especially exciting. As it stands, these processes are still rather inefficient and expensive, rapidly outputting thousands of designs relatively quickly, which translate into meagre yields. **Here we show how the explicit integration of glycosylation as a filter in the 3D *de novo* design pipeline of biologics can significantly improve efficiency and reduce laboratory costs with minimal additional computational resources**. As a proof-of-concept, we used the GlycoShape database and ReGlyco tools (https://glycoshape.org) to filter the results of an open competition launched by Adaptyv Bio in October 2025 for the design of binders as inhibitors against the heavily glycosylated Nipah virus glycoprotein (NiV-G) (https://proteinbase.com/competitions/adaptyv-nipah-competition). Screening of the 1,201 selected designs in block with ReGlyco allows users to eliminate 20% of the confirmed non-binders from the pool in approximately 3 hours on a dual-core CPU. Refinement with a rotamer search enabled flags 11% of non-binders prior to experiment and catches all but 5 confirmed binders from the given predicted 3D complexes. The availability of alternative binding poses improves this score. We complement this analysis with a demo OA colab notebook (https://colab.research.google.com/github/Ojas-Singh/GlycoShape-Resources/blob/main/colab/ReGlyco_Binder_Design_filter.ipynb) to illustrate our workflow. In this demo users can design ‘mini binders’ against human erythropoietin (hEPO) by integrating GlycoShape resources with the RFdiffusion3 (RFD3) pipeline from the Institute for Protein Design (IDP).

## Introduction

Glycosylation is a sophisticated, enzymatically-driven(Narimatsu et al. 2019; Schjoldager et al. 2020) process where glycans are added co-and post-translationally to proteins and processed during trafficking along the secretory pathway. Integration of protein glycosylation in a 3D context is non-trivial because all glycoforms are non-templated and heterogeneous(Fadda et al. 2026; Adams et al. 2025; D. Wu and Robinson 2022; Schachner et al. 2024), both in terms of glycan site occupancy (macroheterogeneity) and type of glycans at each site (microheterogeneity). Glycans come in all shapes and sizes(Varki 2017; Gagneux et al. 2022), which are highly species-specific(Schjoldager et al. 2020; Paschinger and Wilson 2019; De Pourcq et al. 2010; Strasser 2016). Most secreted proteins and biologic therapeutics are glycosylated and the sugars can partially or completely occlude accessible surfaces, delimiting the confines of protein-protein interaction (PPI), guiding multimerization and receptor docking, while affecting folding, stability, trafficking and pharmacokinetic properties of secreted proteins(Egrie and Browne 2001; Tytgat et al. 2019). Current biomolecular *de novo* design and structure prediction protocols are protein-centric(Butcher et al. 2025; Pacesa et al. 2025; Passaro et al. 2025; Abramson et al. 2024; Chai Discovery Team et al. 2025; Ahdritz et al. 2024) and in all approaches currently available post-translational modifications (PTMs), such as glycosylation, are (at best) considered as an afterthought. While we can argue that AI-tools implicitly take into account the effects of PTMs on protein structure due to training(Bagdonas et al. 2021), neglecting to include glycosylation explicitly in a 3D-based protein *de novo* design platform can lead to non-negligible numbers of false positives, needlessly increasing experimental efforts and laboratory costs.

Both AlphaFold 3 (AF3)(Abramson et al. 2024) and RosettaFold All-Atoms (RFAA)(Krishna et al. 2024) allow users to include glycosylation at different stages of the structure prediction process, yet with a very limited repertoire of structures and low accuracy against experiment(Abramson et al. 2024; Krishna et al. 2024). This poor outcome should not be surprising, as the 3D structures of glycans available for AI-training are too few(Fadda et al. 2026), too fragmented and seldom inaccurate(Agirre et al. 2015), despite ongoing efforts in the area(Emsley and Crispin 2018; Dialpuri et al. 2024; Holland et al. 2026). Indeed, data from https://glyconavi.org/ shows that to date only 6,873 proteins in the PDB contain N-glycans, which are the most abundant type of glycans in protein structured domains, for a total of 41,951 glycan structures. Of those 51% consist of a single monosaccharide, with less than 2,054 structures containing 7 or more monosaccharides.

GlycoShape is a structural database and toolbox(Ives et al. 2024) our research team designed to supplement the lack of 3D information on glycans with data obtained from molecular dynamics (MD) equilibrium simulations. To date GlycoShape is the largest and most diverse OA repository of 3D glycan structures worldwide, with over 1,001 unique structures and counting, obtained by clustering analysis of 2 ms of cumulative MD sampling of glycan 3D structures largely from the human glycome, but with examples from all phyla. The 3D database is complemented by ReGlyco (https://glycoshape.org/reglyco), a highly computationally efficient tool that selects glycans structures from the database and links them to the protein at the desired site by minimizing a loss function(Ives et al. 2024).

ReGlyco Ensemble (https://glycoshape.org/ensemble) is a recent expansion of ReGlyco that can be used to select multiple conformers (up to 500) from the weighted ensembles from our in-house MD trajectories(Holborough-Kerkvliet et al. 2026), maintaining the distribution consistency given by the clustering analysis pipeline of the MD data(Ives et al. 2024).

To demonstrate how including glycosylation explicitly with GlycoShape as integral part of the 3D *de novo* design of target-specific binders can increase yields by excluding non-binders from the test pool, and thus reduce laboratory costs, we rescreened the results of the open competition launched in October 2025 by Adaptyv Bio (https://www.adaptyvbio.com/) for the design of binders against the Nipah virus glycoprotein (NiV-G) (https://proteinbase.com/competitions/adaptyv-nipah-competition) as potential inhibitors of viral entry, see **Figure 1**.

**Figure 1.**
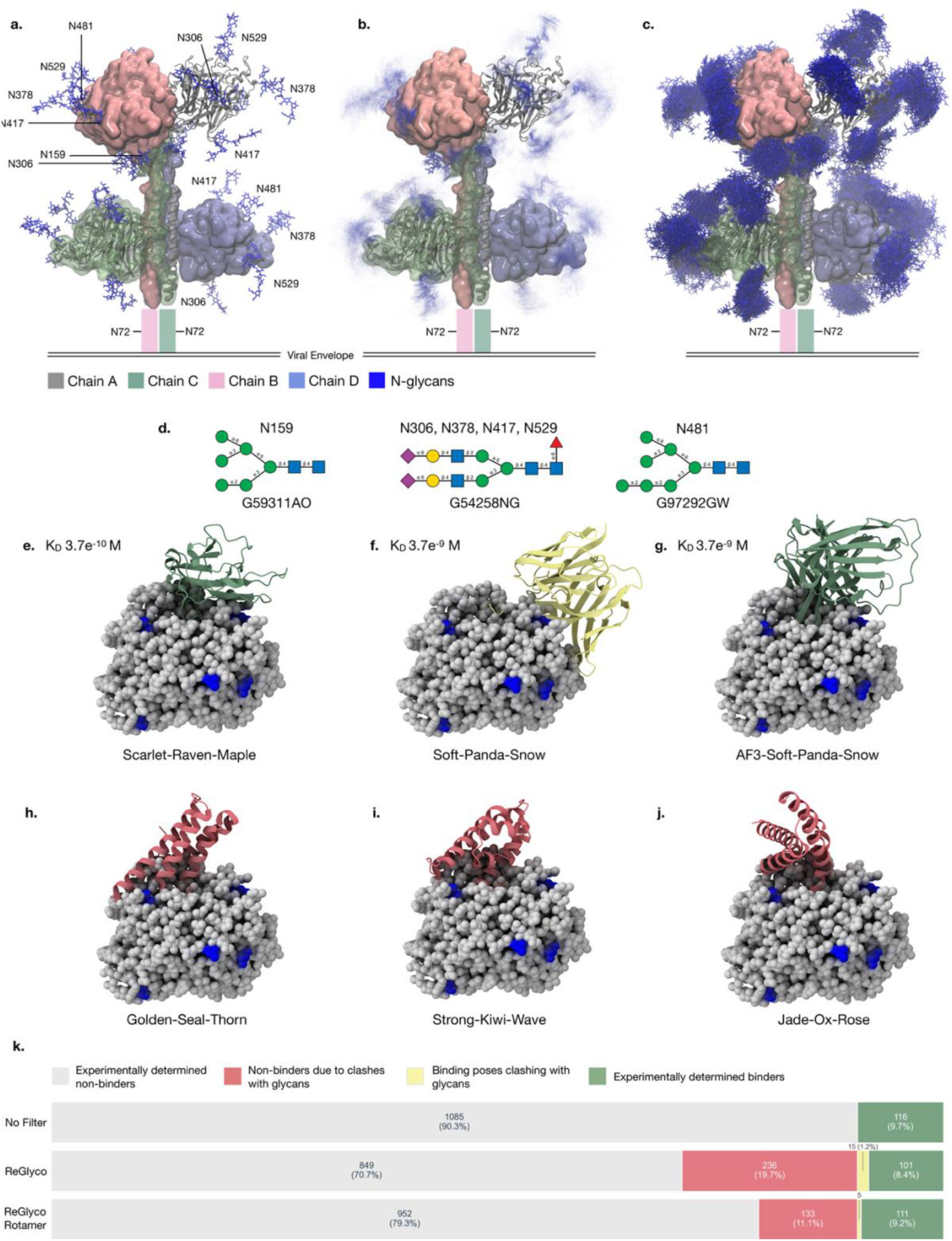
**a.** Structure of the Nipah Virus Glycoprotein (NiV-G) homotetramer (aa 92-602) reconstructed from cryo-EM structures (PDB 7TXZ and 7TY0) bound to broadly neutralising antibody nAH1.3 Fabs(Wang et al. 2022) (not shown). Static (single) glycans 3D structures (shown with sticks in blue) were reconstructed with GlycoShape ReGlyco(Ives et al. 2024). The glycoform selection was guided by glycoproteomics analysis(Hawkins et al. 2025). The visible six glycan sites on chains A, B and D are mapped onto the structure, while glycans on chain C are not labelled for clarity. **b**. Structure of the glycosylated NiV-G reconstructed with GlycoShape ReGlyco Ensemble using 150 frames from the MD trajectory at each site. Rendering effect is obtained with a “long exposure” filter to reflect the glycan dynamics. Sterically constrained glycans and rigid regions of the saccharides can be identified by the darker blue colour. **c**. Structure of the glycosylated NiV-G as in panel b. but with each one of the 150 glycan frames shown in dark blue, to illustrate the potentially occluded volume of the NiV-G protein surface. **d**. Glycans selected at each site represented with SNFG(Neelamegham et al. 2019). Corresponding glycosylation sites are labelled above the SNFG structures and GlyTouCan IDs below. **e-j**. Selected structures from the Adaptyv Bio binder design competition data released (1,201 designs). Proteins are rendered with solid grey spheres and binders with cartoons. Glycosylation sites are shown in blue. Confirmed binders are shown in green cartoons (panels e and g), non-binders in red (h-j), binders but not in the submitted pose, but found to be sterically clashing by ReGlyco, in yellow (f). Binder ids are indicated for all predicted complexes. All non-binders in panels h-j were selected as highest scorers based on a composite score from six structural metrics (*boltz2_complex_iplddt, shape_complimentarity_boltz2_binder_ss, boltz2_min_ipsae, boltz2_ipsae, boltz2_plddt, boltz2_lis*), alongside a binder-like pass count defined as the number of metrics exceeding the median value observed for confirmed binders. All designs shown were successfully expressed in *E. coli*. **k**. Bar chart showing the results of the Adaptyv Bio binder design competition. Results without ReGlyco filtering are shown in the top bar. The middle bar shows the effect of ReGlyco filtering in flagging designs for clashes with one or more glycosylation sites. The bottom bar indicates the effect of the refinement with ReGlyco Rotamer, which introduces flexibility in the Asn sidechain of the aglycon. Molecular rendering with *VMD*(Humphrey et al. 1996) (https://www.ks.uiuc.edu/Research/vmd/) and *pymol* (www.pymol.org).

The Nipah virus (NiV) is a zoonotic henipavirus that can infect humans with an extremely high fatality rate, estimated by the World Health Organization (WHO) at 40% to 75% (https://www.who.int/news-room/fact-sheets/detail/nipah-virus). Outbreaks have been reported periodically since 1998 across south-east Asia, with the most recent one in India in 2026. There are currently no therapeutics or vaccines approved against NiV, while there is an urgent need for the development of strategies to block or to prevent infection. NiV infection requires fusion of the viral envelope with the host cell membrane. This process is mediated by the concerted action of two envelope glycoproteins, namely NiV-G(Wang et al. 2022) and the NiV fusion (NiV-F) glycoprotein(Xu et al. 2015). NiV-G initiates entry by binding to the transmembrane protein tyrosine kinases ephrin-B2 or ephrin-B3 as receptors(Xu et al. 2008; Bonaparte et al. 2005; Bowden et al. 2008; Negrete et al. 2005).

In Oct 2025 Adaptyv Bio (https://www.adaptyvbio.com/) launched an open competition for the *de novo* design of binders targeting NiV-G as inhibitors of ephrin-B2 interaction, and thus based on the crystal structure of the NiV-G monomer in complex with ephrin-B2(Bowden et al. 2008) (PDB 2VSM). Of 10,000+ submissions of novel ephrin-B2-based binders, with entries within a range between 100 and 150 kDa, 1,201 designs were selected for expression in *E. coli*, with 1,028 successfully expressed (86%). Only 116 of the submitted entries selected were confirmed as binding NiV-G (10% success rate), see **Figure 1.k**. None of the entries included glycosylation explicitly of either the binder or of the target NiV-G. While the binders were expressed in *E. coli* for the binding assays, and thus were not glycosylated, the NiV-G was expressed in HEK293 cells and thus it was fully glycosylated, as shown in **Figures 1.a-c**.

To illustrate the extent of the glycosylation of the NiV-G targeted for design here, we reconstructed the glycans with ReGlyco based on the NiV-G cryo-EM structure(Wang et al. 2022) and produced a representative fully glycosylated tetramer based on recently published glycoproteomics data(Hawkins et al. 2025), see **Figure 1.a-c**. We used ReGlyco Ensemble to link the glycan structures to the protein scaffold using 150 frames from the MD trajectories for each chosen glycan, see **Figures 1.b** and **1.c**. As a note of interest, the rendering in **Figure 1.b** is obtained with a bespoke filter we designed to mimic the effect of a long-exposure photograph. This perspective highlights the glycans that are less dynamic, and thus better resolved, because of inherent structural restraints, or protein surface complementarity, while blurring out the contributions from the most dynamic fragments, providing a potentially more realistic graphical representation of glycan shielding and screening relative to the timeframe overlay shown in **Figure 1.c**. The results of our filtering are presented and discussed below. As a complement to this case, we present an OA colab notebook demo to illustrate our workflow where users can run a 3D *de novo* design of mini-binders to the heavily glycosylated hEPO by integrating GlycoShape resources with RFdiffusion 3 (RF3) protein design pipeline(Butcher et al. 2025).

### Filtering the NiV-G binding competition designs with ReGlyco and ReGlyco Rotamer

We downloaded and screened the 1,201 designs selected to date for expression by Adaptyv Bio to filter potential clashes due to the NiV-G extensive glycosylation, see **Figure 1.a-c**. The 3D structures of all binders sequences submitted were predicted with Boltz-2(Passaro et al. 2025) for scoring and selection purposes. We used those 3D structures in our filtering pipeline. We first used the GlcNAc scanning mode(Ives et al. 2024), currently set as the ‘default’ option in the reconstruction pipeline on ReGlyco (https://glycoshape.org/reglyco), to identify the glycosylation sites on the structure of the NiV-G monomer structure (PDB 2VSM), which was used as a 3D model of the receptor in the open competition. The GlycoShape GlcNAc scanning(Ives et al. 2024) tool correctly identified six glycosylation sites(Hawkins et al. 2025) as potentially occupied, namely N159, N306, N378, N417, N481 and N529. We linked to all of these a Man_5_-GlcNAc_2_-oligomannose (GlyTouCan ID G000208MO) for screening purposes with two filtering approaches, 1) ReGlyco, where the glycan is linked to the aglycon Asn sidechains in the same conformation as the one predicted by Boltz-2, i.e. rigid rotamers, 2) ReGlyco Rotamer, a new tool we developed for this work, and available on ReGlyco ‘Advanced (site-by-site) Glycosylation’ mode, that allows the aglycon Asn sidechain torsions to access different rotamers based on Dunbrack’s Smooth Backbone Dependent Rotameric Library(Shapovalov and Dunbrack 2011) (https://dunbrack.fccc.edu/lab/bbdep2010) to clear steric clashes that depend on the Asn sidechain orientation alone(Ives et al. 2024).

The results of the competition show that only 116 designs out of the 1,201 that Adaptyv Bio selected for protein expression in *E coli* were found positive for binding to NiV-G expressed in HEK293 cells, see **Figure 1.k**; 173 were not successfully expressed, and 917 did not bind NiV-G. Parsing the 1,201 binders designs in-block through the ReGlyco filter took 3 hours and 16 minutes on a dual core CPU. ReGlyco flagged 251 complexes as non-binders due to steric clashes involving at least one glycosylation site. Of the 251 designs flagged, 236 were among the confirmed non-binders by experiment and 15 were among the confirmed binders, see **Figure 1.k**. Refining the screening by enabling rotameric freedom through ReGlyco Rotamer reduced the number of 251 designs flagged by Re-Glyco for steric clashes to 138. Of those 133 were among the confirmed non-binders and 5 among the confirmed binders, see **Figure 1.k**. We visually assessed the clashes in these 5 entries and decided to rebuild the corresponding complexes with the AF3 server (https://alphafoldserver.com/) with no added glycans, to check for binding poses alternative to Boltz-2 predictions provided online(Overath et al. 2025). In all but one of those five cases, namely the ‘Soft Panda Snow’ (K_D_ = 4 μM) entry shown in two poses, the Boltz-2 one and the AF3, in **Figures 1.f** and **1.g**, respectively, AF3 produced a non-clashing pose. This result suggests that the use of different protein structure and complexes prediction algorithms may also help to refine the pool of potential binder designs to promote to experimental testing. All complexes from the Adaptyv Bio Re-Glyco filtering analysis are available on https://glycoshape.org/downloads/adaptyv-nipah-competition.

As a note of interest, our first attempt to build all 5 binders flagged by ReGlyco on the AF3 server with a fully glycosylated NiV-G monomer as a target, timed-out with no output after 24 hrs. We re-run those attempts more recently and each prediction took 4 hrs to terminate successfully. This indicates that the AF3 algorithm is continuously improving as a very useful prediction tool for complexes involving glycosylated biomolecules, yet extremely time consuming (to date) relative to the AI+ReGlyco scheme we present here and discuss below within the framework of an OA demo platform for mini-binders design.

### A colab notebook for the design of binders to hEPO with RFDiffusion3 and ReGlyco

Human erythropoietin (hEPO) is a 30 kDa glycoprotein hormone that regulates the production of red-blood cells, or erythropoiesis, by binding a transmembrane dimeric receptor(Syed et al. 1998) (hEPOR), see **Figure 2**, located on the membrane of erythrocytes. hEPO is a heavily glycosylated biologic bearing three fully occupied N-glycosylation sites at N24, N38 and N83, and one O-glycosylation site at S126(S. Wu et al. 2025; Murakami et al. 2016). Glycosylation, and more specifically sialylation, is a key determinant of hEPO pharmacokinetic properties, so that the hyperglycosylation of hEPO has been engineered successfully to produce a recombinant form of the hormone, known as NESP(Egrie and Browne 2001; S. Wu et al. 2025), with remarkably higher half-life. The extensive degree of hEPO glycosylation, and the position of the two additional sites in NESP, lead the hormone to bind hEPOR in designated (non-glycosylated) surface areas, see **Figure 2.b**. Here we illustrate our ReGlyco Binder Design Filter workflow through a demo colab notebook using hEPO as a sample target receptor. This choice is due primarily to hEPO low molecular weight compared to viral fusion proteins such as NiV-G and NiV-F discussed earlier, which are incompatible with the 16 GB GPU limit of the free notebook.

**Figure 2.**
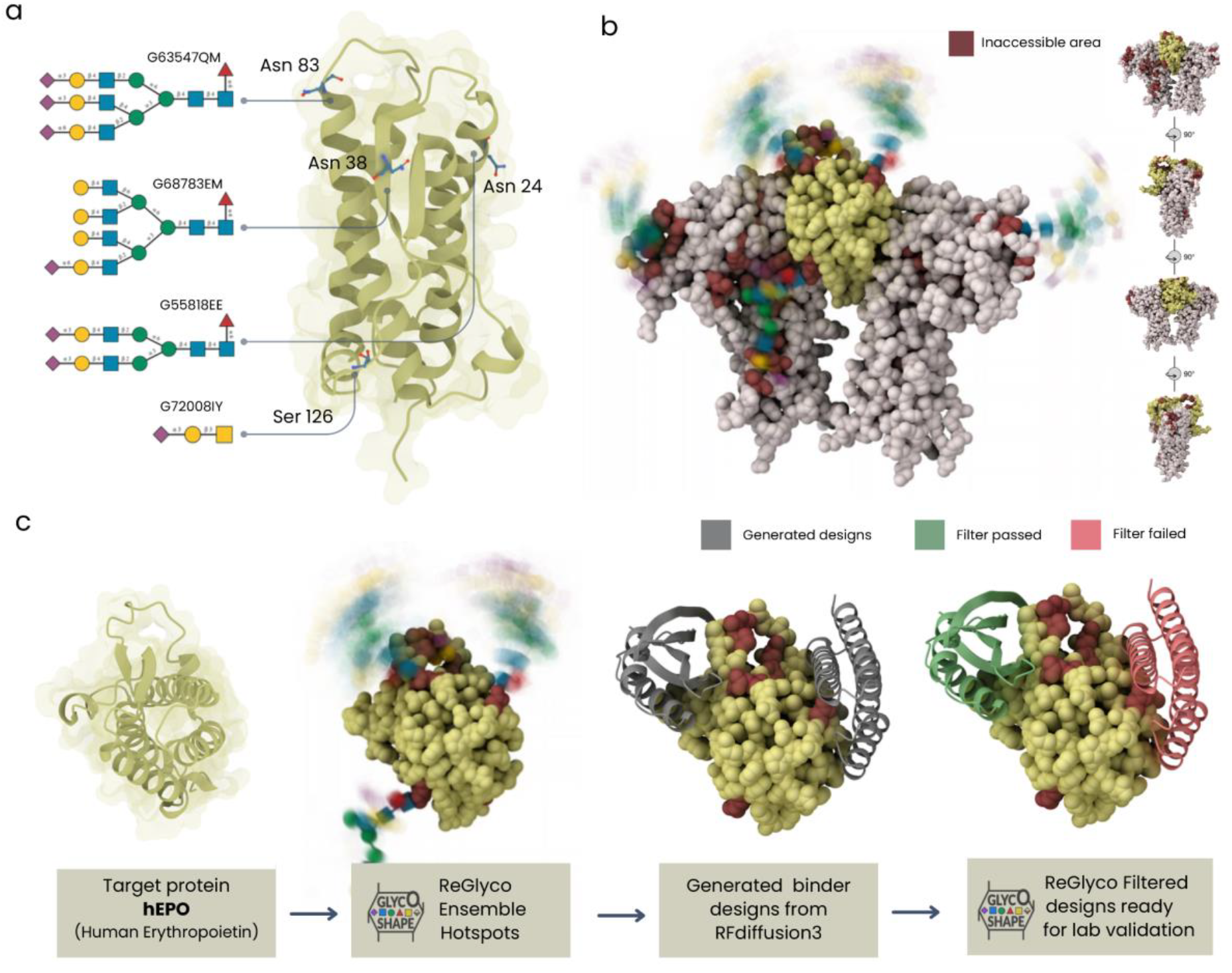
**a.** 3D structure of the human erythropoietin (hEPO) (from PDB 1EER) rendered in yellow cartoons with a transparent molecular surface. N-linked (3 sites) and O-linked (1 site) glycan sites are labelled according to the PDB numbering. The glycans selected for the protein reconstruction are shown in SNFG with GlyTouCan IDs. **b**. 3D structure of hEPO (yellow spheres) bound to the hEPO receptor (white spheres) (PDB 1EER). The exclusion surface areas due to glycosylation are highlighted with dark red spheres. Glycans are rendered in SNFG with the ‘long exposure filter’ to show their dynamics modulated by the protein surface at and around the site. On the right side of the panel the structure on the complex is progressively rotated over 360° in 90° steps around the z axis. **c**. Vignettes showing in four stages the RF3-ReGlyco mini-binder design workflow to hEPO as in the demo (https://colab.research.google.com/github/Ojas-Singh/GlycoShape-Resources/blob/main/colab/ReGlyco_Binder_Design_filter.ipynb). From the left-hand side, the 3D structure of hEPO is selected (yellow cartoons with a transparent molecular surface) as target protein. ReGlyco Ensemble is used to reconstruct the glycoprotein with the desired glycosylation. This process will output the inaccessible surface area due to the presence of glycans (dark red spheres) and the accessible surface area (yellow spheres) that can be targeted for binding, or ‘hotspots’. RF3 is used to design mini-binders. Generated designs are shown in grey cartoons. These designs are filtered by ReGlyco to exclude designs clashing with one or more glycosylation sites: predicted binders are shown in green cartoons (filter passed) and predicted non-binders (filter failed passed) in red cartoons. Mol* (https://molstar.org/) was used for structure rendering.

Additionally, the glycoengineering of recombinant hormones and of small biologics is also of high interest(Madsen et al. 2020; Bousfield and Harvey 2019; Flintegaard et al. 2010), as well as the promise and ambitions of *de novo* design technologies in this area.

The ReGlyco Binder Design Filter demo workflow is illustrated in **Figure 2.c** in four vignettes and explained step-by-step below. In the default mode (‘quick’ demo mode), the target receptor is hEPO (from PDB 1EER) with glycosylation selected at the known sites(S. Wu et al. 2025; Murakami et al. 2016) and glycan types, indicated by GlyTouCan(Tiemeyer et al. 2017) IDs, aligned with published glycomics profiles, see **Figure 2.a**. Users can leave the defaults for the bundled hEPO example, or switch to ‘advanced’ mode to upload a different target, provide their own glycan list, or change how the interaction patch is selected. Details on how to set up the ‘advanced’ mode are in the notebook, at the bottom of the workflow.

We set a limit of up 250 aa for the total length of the complex due to restrictions in the free version of the colab notebook resources. The user can select up to 6 residues as interaction hotspots and up to 20 mini-binders as outputs across all available hotspots regions. Each mini-binder is set up by default to a length of 80 aa, but this parameter can be changed to up to 130 aa. Three options are available to distribute the binders across the selected hotspots:

1) ‘chunked’ (default) to spread binders across different hotspots in the pool, 2) ‘random’ to randomly select different hotspots for each binder, and 3) ‘rotate’ to select hotspots in order as a target for each binder. Re-Glyco Ensemble is run to identify the surface areas of the receptor inaccessible due to the specific glycosylation selected, i.e. the ‘glycosylation hotspots’. The output can be inspected visually in the Mol* GUI(Sehnal et al. 2021). At this point the demo version will generate a subset of 20 mini-binders using RFDiffusion3(Butcher et al. 2025) (RFD3) resources against the protein-only target. Each design candidate is set-up to target a different hotspot to produce a broad spread of designs that ReGlyco can triage downstream. The results can be viewed in the Mol* GUI(Sehnal et al. 2021). The designs produced are then filtered by ReGlyco for clashes with one or more glycosylation sites. This test can result in three outcomes, 1) ‘pass’, when no clashes are detected, 2) ‘borderline’, clashes are cleared by enabling rotameric degrees of freedom through ReGlyco Rotamer, and 3) ‘fail’ clash persists indicating serious overlap with one or more glycosylation sites. The demo’s default settings lead to 14 designs in the ‘pass’ category, 1 in the ‘borderline’, and 5 in the ‘fail’ category due to irresolvable clashes with one or more glycosylation sites. All results with their respective classification label, can be viewed in the Mol* GUI window (Sehnal et al. 2021).

## Conclusions

In this work we have demonstrated that glycosylation can significantly limit the spatial accessibility of the target receptor’s protein surface, directly affecting yields in *de novo* design processes. As a case study, our screening of 2025 Adaptyv Bio open competition’s results for the design of inhibitors to the NiV entry, highlights that ReGlyco and ReGlyco Rotamer can efficiently flag non-binders prior to experimental testing at zero-cost. We suggest that the filtering should be implemented as an entry optimization step by user submitting designs and/or as to refine the pool of candidates progressing to experimental testing, or both. We filtered all 1,201 entries with both ReGlyco and ReGlyco Rotamer and showed that this simple additional step can reduce laboratory effort and costs with minimal additional computing resources. ReGlyco can successfully screen 1,201 complexes in approximately 3 hrs time on a dual-core CPU, flagging 20% (236) of non-binders due to steric clashes with glycans. Standard implementation of ReGlyco also flags as clashing 1.2% of actual binders. Refinement with ReGlyco Rotamer reduces the number of non-binders flagged to 11% (133) but clears most of the actual binders, with only 0.4% (5) flagged as clashing. A careful analysis of those 5 complexes confirms that the clash is irresolvable for the given complex prediction and for that reason the binding pose may be incorrect. Indeed, we have shown that changing the prediction algorithm provides alternative (non-clashing) poses in at least one case, suggesting that the use of multiple structure prediction tools could also improve the efficiency of the *de novo* design pipeline.

## Acknowledgements

Research Ireland/Taighde Éireann (former SFI) Frontiers for the Future Programme is gratefully acknowledged for financial support of OS postgraduate training (20/FFP-P/8809). UKRI BBSRC (UKRI2974) is gratefully acknowledged for funding OS post-doctoral training at the University of Southampton. EF and OS acknowledge IT Solutions at the University of Southampton for generous allocation of computational resources on the IRIDIS High Performance Computing Facility, and the ORACLE Cloud Infrastructure (OCI) team, in particular Diarmuid McKenna and Derek Cullen, for their continued, expert support and for the seamless hosting of the GlycoShape server and applications on OCI. EF and OS are grateful to Harjasnoor Kakkar for her time and for insightful discussions and advice on the matters discussed in this manuscript.

## Data Availability

Details on the ReGlyco Binder Design Filter pipeline and algorithms are provided in Supplementary Material. The ReGlyco Binder Design Filter colab notebook is available at https://colab.research.google.com/github/Ojas-Singh/GlycoShape-Resources/blob/main/colab/ReGlyco_Binder_Design_filter.ipynb The GlycoShape database and tools used in the colab notebook are available at https://glycoshape.org. The results of the ReGlyco filtering of the Adaptyv Bio Open Competition are available on https://glycoshape.org/downloads/adaptyv-nipah-competition.

