## Supplementary Information for "Integrating glycosylation in *de novo* protein design with ReGlyco Binder Design Filter" (from another version of this preprint: version 1, 2026-04-17)

**Methods**

**ReGlyco glycoprotein builder**

To model glycosylated variants of a protein, we first defined the user-selected or GlcNAc scan^1^ predicted glycosylation sites $S=\{1,\ldots,m\}$ on the bare protein structure $P=\{p_{i}{\}}_{i=1}^{N_{P}}$. For each site $s\in S$, ReGlyco^1^ assigned a glycan-specific conformational library $L_{s}=\{G_{s,k}{\}}_{k=1}^{K_{s}}$. Here, $G_{s,k}$ denotes a representative glycan structural cluster, which was attached to the acceptor residue using a rigid-body map $T_{s}$ that aligned the glycan linkage atoms with the local protein geometry at site $s$. Linkage torsions $\left( \phi_{s},\psi_{s},\theta_{s} \right)$ were then sampled from glycan-specific circular priors derived from molecular dynamics simulations or Privateer^2^.

A candidate glycoprotein state was defined by $z_{s}=\left( k_{s},\phi_{s},\psi_{s},\theta_{s} \right)$, and the fully assembled structure was written as

$$X\left( z \right)=P\cup{\bigcup^{m}}_{s=1}T_{s}\left( k_{s},\phi_{s},\psi_{s},\theta_{s} \right) G_{s,k_{s}}.$$

We evaluated each sampled configuration using a steric compatibility score, calculated as $S\left( X \right)=\sum_{\left( a,b \right)\in C\left( X \right)} \rho\left( d_{ab} \right)$, where $C\left( X \right)$ denotes potentially clashing atom pairs across protein-glycan and glycan-glycan contacts, $d_{ab}$ is the interatomic distance, and $\rho\left( \cdot\right)$ is a short-range overlap penalty. ReGlyco filtered the sampled states to retain only sterically viable conformations satisfying

$$S\left( X\left( z \right) \right)\leq\tau,$$

where $\tau$ is the steric acceptance threshold. Ultimately, this process combines library selection, torsion sampling, and steric filtering to generate a re-glycosylated structure consistent with both local protein geometry and glycan conformational priors.

**Rotamer enabled sampling**

For glycosylation sites exhibiting local side-chain flexibility, we expanded the sampled state space to include a residue rotamer variable. For this purpose, we developed ReGlyco Rotamer, which enables the aglycon Asn side-chain torsions to sample alternative rotamers from Dunbrack’s Smooth Backbone-Dependent Rotameric Library according to the local backbone dihedrals $\left( \Phi_{s},\Psi_{s} \right)$.This higher sophisticated filtering mode ensures clearance of steric clashes arising from the Asn side-chain orientation alone. Each sampled rotamer reoriented the acceptor side chain prior to glycan placement, thereby modifying the local attachment geometry available to the glycan.

The combined state of site $s$ became $\left( k_{s},\phi_{s},\psi_{s},\theta_{s},r_{s} \right)$, and the algorithm accepted configurations satisfying the steric criterion $S\left( X\left( z,r \right) \right)\leq\tau$. This extension allowed the method to resolve steric conflicts not only by repositioning the glycan, but also by actively reorienting the acceptor side chain.

**ReGlyco Ensemble**

To accurately capture the conformational diversity of glycans, ReGlyco Ensemble^3^ repeated the sampling-and-filtering procedure to retain a comprehensive collection of accepted configurations,

$$E=\{X^{\left( n \right)}{\}}_{n=1}^{N}.$$

Each member of this ensemble corresponds to a sampled set of variables, $z^{\left( n \right)}$, that satisfies the steric acceptance condition. Furthermore, if multiple protein backbone models $P^{\left( b \right)}$ were available, sampling was performed across both backbone and glycan states,

$$X^{\left( n \right)}=X\left( P^{\left( b_{n} \right)},z^{\left( n \right)} \right).$$

This approach successfully propagated both protein and glycan conformational variability into the final ensemble. This multi-model structure therefore represents a feasible conformational ensemble rather than a single optimized state, allowing us to summarize site-specific glycan behaviors, such as the observed frequency of clusters and ensemble-averaged torsions.

**Hotspots calculation from glycoprotein ensemble**

We quantified the degree of glycan shielding by comparing solvent accessibility before and after glycosylation across the generated ensemble. For each protein residue $i$, we calculated the solvent accessible surface area (SASA) of the unglycosylated bare protein, $A_{i}^{bare}$, and the corresponding SASA in each glycosylated ensemble member, $A_{i}^{\left( n \right)}$. The glycosylated ensemble-average SASA was

$$A_{i}^{glyco}=\frac{1}{N}\sum_{n=1}^{N} A_{i}^{\left( n \right)}.$$

The shielding, or coverage fraction, induced by the glycans was defined as

$$C_{i}=\frac{A_{i}^{bare}-A_{i}^{glyco}}{A_{i}^{bare}}.$$

Using these metrics, we defined a binary hotspot mask $H_{i}$ to identify residues that remained exposed despite the surrounding glycan shielding. We set $H_{i}=1$ only when a residue met three strict criteria: it was surface-exposed in the bare protein ($A_{i}^{bare}>0$), it remained accessible in the glycosylated ensemble ($A_{i}^{glyco}>\varepsilon$), and it was not shielded by more than $30\%$ on average ($C_{i}\leq0.30$). Otherwise, we set $H_{i}=0$. This resulting binary mask served as a robust, residue-level epitope accessibility map for downstream binder-design analysis.
